# From Photoperiod Thresholds to Photoperiod sensitivity: Dual Strategies for Cost-Effective Speed Breeding and Climate-Ready Barley

**DOI:** 10.1101/2025.09.04.674238

**Authors:** Nicola Rossi, Wayne Powell, Karen Halliday, Rajiv Sharma

## Abstract

Yield and the duration of the growing season are closely linked. Climate change may shorten growing seasons in certain European regions, and reducing the time to flowering could be an effective strategy to mitigate its effects. Therefore, exploring allelic combinations shape flowering time, is needed. Additionally, speed breeding (SB), characterized by extended photoperiods to accelerate generation time, can be energy-intensive, and the shortest day length needed to induce rapid flowering remains unknown.

We present the first integrated study of how allelic variation at three key flowering time genes, *PPD-H1, ELF3* and *PHYC*, modulates three parameters of the photoperiod response model: threshold photoperiod, photoperiod sensitivity, and intrinsic earliness. We recorded flowering under lengths of 16–24h in Near Isogenic Lines carrying *PhyC-e* or *PhyC-I* allele within *ppd-H1* background, and in lines from HEB-25 combining wild and domesticated alleles of *ELF3* and *PPD-H1*. The *ELF3* allele in *ppd-H1* background reduced intrinsic earliness, whereas *PhyC-e* reduced photoperiod sensitivity, opening opportunities for climate change adaptation. Remarkably, *ppd-H1* lines flowered at a 20-h threshold, whereas *Ppd-H1* lines showed no response, consequently we propose new SB photoperiods at 20 and 16h depending on *PPD-H1* background. These photoperiods lower energy costs compared to the current 22h standard.

**Highlights:** By studying barley key photoperiod response genes, our results support energy-efficient speed breeding and the development of climate-resilient varieties through targeted genetic control of flowering time.

## Introduction

In cereal crops, flowering time is a crucial trait for both adaptation (Jones et al., 2008; Faure et al., 2012) and for optimizing potential yield (Wang et al., 2009;Mäkinen et al., 2018;), as these factors are often closely linked (Slafer, et al., 2023). Flowering time is also an important consideration in speed breeding, which aims to shorten generation time (Hickey et al., 2017; Watson et al., 2018; Chiurugwi et al., 2019; Cazzola et al., 2020; Mobini et al., 2020; Samineni et al., 2020; Fang et al., 2021; Schilling et al., 2023). According to the breeder’s equation (Lush, 1937), genetic gain per unit time can be increased not only by applying selection but also by reducing generation time. Shortening the generation cycle, as in speed breeding, accelerates the rate at which homozygosity is achieved, thereby facilitating the fixation of alleles and potentially increasing the accuracy of selection and narrow-sense heritability in subsequent generations. The ability of crop plants to respond to increasing day lengths in controlled environments is instrumental in refining protocols that accelerate generation turnover, thereby enhancing genetic gain (Cobb et al., 2019). Speed breeding protocols manipulate conditions like photoperiod, temperature and light intensity to enable multiple generations per year. For long day plants (LDP) such as barley, wheat, canola, chickpea, pea and oat, speed breeding protocols typically employ a 22-hour photoperiod combined with a 2-hour dark phase and cooler night temperatures to accelerate generation turnover (Watson et al., 2018;González-Barrios et al., 2021). The 2-hour dark period, alongside reduced night temperatures, facilitates plant recovery and minimizes stress linked to accelerated growth (Watson, A., 2019). However, this approach may overlook key findings from photoperiod response modelling in long-day plants (Major, 1980) and has limitations when diverse germplasm (e.g. wild relatives) is used (Rossi et al., 2024). Experiments investigating major flowering time genes under defined photoperiod conditions have enhanced our understanding of the genetic control of flowering, informing strategies to improve adaptation in breeding programs (Pérez-Gianmarco et al., 2019). This depth of understanding presents a strategic opportunity: by targeting photoperiod-responsive genes within a highly isogenic background, we can more precisely investigate how extended photoperiods affect flowering time—advancing our understanding of adaptation, informing breeding decisions, and refining speed breeding protocols. Among genetic factors, flowering time in small-grain cereals such as wheat and barley is governed by genotype and genotype-by-environment interactions, with the photoperiod and vernalization pathways serving as central regulators of day-length and low-temperature responses, respectively. In barley broad adaptation arises from extensive allelic variation within these pathways (Fernández-Calleja et al., 2021), which distinguishes winter and spring growth habits. Spring barley genotypes exhibit null or reduced vernalization requirements, relying predominantly on photoperiod sensitivity to delay flowering under long days. Once photoperiod and vernalization responses are saturated, residual genotypic variation in flowering time is mainly controlled by earliness per se (eps) loci (Slafer et al., 1994; Parrado et al., 2023).

In barley (Hordeum vulgare L.), flowering time exhibits a characteristic bi-linear response to photoperiod duration. The seminal work by Major (1980), in relation to the photoperiodic response, established that small grain cereals display: (i) an initial linear decline in flowering time with increasing photoperiod until reaching a threshold (*threshold photoperiod*), beyond which (ii) daylength extension no further accelerates development. The slope of the initial linear phase (*photoperiod sensitivity*) quantifies the responsiveness to daylength, while the stable phase represents the degree of *intrinsic earliness* - the minimum flowering time when photoperiod requirements are fully satisfied, such parameter is known to be controlled by earliness per se genes such as barley *CENTRORADIALIS* (*CEN*) (Fernández-Calleja et al., 2021). In barley and other temperate cereals, key genes such as wheat *PHOTOPERIOD1 (PPD1)* and its barley orthologue *PPD-H1*, as well as earliness per se genes, underpin variation in these photoperiodic traits. Pérez-Gianmarco et al. (2019) applied Major’s (1980) photoperiod response model to characterize *PPD1* alleles in wheat (*Triticum aestivum* L.), revealing conserved threshold photoperiod and intrinsic earliness across genotypes, with variation occurring primarily in photoperiod sensitivity. In barley (*Hordeum vulgare* L.), Parrado et al. (2023) conducted complementary studies using near-isogenic lines (NILs) for *PPD-H1* across controlled and field environments. While their study did not employ formal modelling, the results demonstrated that *PPD-H1* alleles modulate both photoperiod threshold and sensitivity, while maintaining stable intrinsic earliness.

Photoperiodic flowering in grasses is regulated by a network of genes that coordinate environmental signals with developmental responses. Central to this network is the gene *PPD1*, which acts as a molecular switch to initiate flowering in response to day length (Turner et al., 2005). *PPD1* encodes a pseudo-response regulator protein (*PRR37*) that promotes the expression of *FLOWERING LOCUS T1* (*FT1*), the key integrator of flowering signals, leading to floral transition under long days (Shaw et al., 2020). The regulation of *PPD1* expression results from the interplay between the circadian clock and light signalling pathways, such that its activation occurs when the timing of gene expression coincides with periods of light under long-day conditions (Song et al., 2015). This mechanism ensures that *PPD1* is only activated when internal circadian rhythms align with specific external cues, such as light. Upstream regulators of *PPD1* include the circadian clock component *EARLY FLOWERING 3* (*ELF3*) and light-sensing phytochromes, especially *PHYTOCHROME C (PHYC)*. In model grasses such as *Brachypodium, PHYC* has been shown to likely repress *ELF3* post-translationally (Bouché et al., 2022; Alvarez et al., 2023; Gao et al., 2023). As a result, *ELF3* cannot suppress *PPD1*, leading to the flowering response via the induction of *FT1* expression. However, the precise molecular interactions among these clock components in barley are less well characterized, and much of our current understanding is extrapolated from these related grass (*Brachypodium*, wheat and rice) systems. Nevertheless, research in barley has confirmed the roles of functional allelic variation at *PHYC, PPD-H1* (*PPD1* orthologue) and *ELF3* in modulating photoperiod sensitivity and flowering time (summarized below).

The effects of variation at *PPD-H1* indicate the presence of two functional alleles, the dominant wild allele *Ppd-H1*, which leads to early flowering phenotypes and the recessive *ppd-H1* harbouring mutation within the CCT domain which is thought to diminish its ability to activate *FT1* compared to *Ppd-H1* (Turner et al., 2005) and leads to a delay in flowering. The recessive mutation giving rise to *ppd-H1* favoured the expansion of barley from the Fertile Crescent to higher latitudes (von Bothmer et al., 2010), characterized by longer growing seasons. Therefore, *ppd-H1* is preferred in regions characterized by long growing seasons (such as central and northern Europe) and *Ppd-H1* in environments characterized by higher temperatures and drought (e.g. the Mediterranean basin) (Wiegmann et al., 2019). The *ELF3* allelic series comprises three main alleles: the domesticated allele *Elf3*, the *elf3* alleles, and the wild *ELF3*_*Hsp*_ found in *Hordeum spontaneum* lines. The *elf3* alleles comprise two alleles, the *eam8*.*k* which contains two deletions, one inversion, and two small insertions and *eam8*.*w* allele which has a point mutation that causes a premature stop codon (Faure et al., 2012; Zakhrabekova et al., 2012; Zahn et al., 2023). These mutations result in photoperiod insensitivity and early flowering both in long and short days, likely due to lack of repression of *PPD-H1*, which enhances *FT1* activation and disrupts the circadian clock (Müller et al., 2020). The wild *ELF3*_*Hsp*_ allele is thought to contain a non-synonymous mutation at amino acid position 669, contributing to an acceleration of flowering (Zahn et al., 2023). *elf3* alleles have been recognized as a crucial factor aiding barley’s adaptation to very short growing seasons at high latitudes (Faure et al., 2012). Allelic series at barley’s *PHYC* gene involve the wild *PhyC-I* and the *PhyC-e* allele that harbours a mutation in a critical position within the GAF domain, located at the end of a helix near the chromophore pocket. This mutation causes a notable reduction in flowering time. *PhyC-e* is thought to bypass the circadian clock genes (of which the Evening Clock-a circadian clock regulatory complex - belongs to), inducing *PPD-H1* in barley which then leads to an enhanced accumulation of *FT1* and early flowering (Nishida et al., 2013; Pankin et al., 2014).

The importance of *ppd-H1*, which contributes to extending the growing season at moderately long days, may decline in some European regions as temperatures continue to rise. It has been suggested that *Ppd-H1* could play a more prominent role under these conditions (Herzig et al., 2018), given its strong effect on accelerating flowering. However, relying solely on *Ppd-H1* may not always offer the optimal balance for adaptation. This highlights the need to explore allelic combinations in a *ppd-H1* background that support intermediate flowering times—offering greater flexibility for adapting to warmer climates without excessively shortening the growing season. The effect of *ELF3*_*Hsp*_ and *Phyc-e* in a *ppd-H1* background could lead to intermediate phenotypes that shorten the growing season enough to escape terminal heat and drought associated with the increase in temperatures, but still guaranteeing a longer growing season than the one achievable with the effect of *Ppd-H1*.

A recent study by Rossi et al. (2024) demonstrated that *ELF3* and *PPD-H1* are key regulators of developmental timing under both standard (i.e. 16 hours) and speed breeding (i.e. 22 hours) photoperiods. Notably, this revealed that domesticated alleles benefit the most in accelerating the growing cycle under speed breeding conditions. These findings highlighted that the effectiveness of speed breeding protocols is highly influenced by allelic variation, particularly within diverse germplasm pools.

However, the photoperiod threshold required to trigger accelerated development—and its interaction with genotype—remains unexplored, posing a key limitation to the design of efficient and energy-smart speed breeding protocols. Cutting energy could reduce lighting-related energy costs by approximately 4.54% each hour. To put these savings into perspective, Zhang et al., (2017) reported that producing 653 kg of tomatoes over 15 years from an 8 sq. ft. area required a total 29,250 kWh with 600-W LED lights. This equates to around 45 kWh/kg/year for LED lighting. For vertical farming, the average is approximately 250 kWh/kg/year (Teo et al., 2024). In commercial breeding facilities, reducing LED lighting by one hour per day can save about 2.3 kWh annually per 8 sq. ft. These savings scale up in larger operations, illustrating how genotype-informed speed breeding protocols can make breeding programs both more sustainable and cost-effective.

Our study builds upon our recent study (Rossi et al. (2024))on investigating how wild and domesticated alleles of *ELF3* and *PPD-H1* from the Nested Associated Mapping population HEB-25 (Maurer et al., 2015), as well as *PHYC* (alleles *PhyC-e, PhyC-I*) assessed using Bowman introgression lines (Druka et al., 2011), influence flowering time under varying photoperiods (16–24h). By integrating phenotypic data with gene expression profiles, we seek to gain insight into the development of climate-resilient barley varieties by characterising the contribution of key alleles to photoperiod sensitivity and intrinsic earliness. Additionally, we determine the photoperiod saturation threshold to optimize energy-efficient speed breeding.

## Materials and methods

### Plant material

In this study, we investigated the photoperiod response model in two genetically distinct plant groups. The first group, referred to as the “HEB group,” consists of four recombinant inbred lines (RILs) from a HEB-25 family of the multiparent nested associated mapping (NAM) population “Halle Wild Barley” (HEB-25). This population was created by crossing 25 wild barley parents—24 *Hordeum vulgare* ssp. *spontaneum* (Hsp) and one *Hordeum vulgare* ssp. *agriocrithon*—with the spring barley cultivar Barke (*H. vulgare* ssp. *vulgare*, Hv). The resulting progeny were backcrossed to the female elite barley variety Barke, followed by three generations of selfing through single-seed descent (BC1S3), and further propagated minimum to the BC_1_S_6_ generation. More details about the population development is given in Maurer et al., (2015).

The HEB group includes the four possible allele combinations at the *ELF3* and *PPD-H1* loci. To ensure the most isogenic background possible available, these combinations were identified based on the lack of segregation at markers linked to four key flowering time genes (*CEN, SDW1, VRN-H1/PHYC*, and *PPD1*), as determined using the Infinium iSelect 50k SNP array (Maurer & Pillen, 2019). The selected markers associated with the target flowering time genes were chosen based on the subset used for the HIF pre-selection in Zahn et al. (2023), the markers composition is provided in Table S1.

In the HEB group we designate the alleles as follows: the dominant *Ppd-H1* as wild *PPD-H1*_*Hsp*_, the recessive *ppd-H1* as domesticated *PPD-H1*_*Hv*_, the domesticated *Elf3* as *ELF3*_*Hv*_ and the wild *ELF3*_*Hsp*_. Consequently, the factorial combination of *ELF3* and *PPD-H1* includes four lines: *ELF3*_Hv_/*PPD-H1*_Hv_, *ELF3*_Hv_/*PPD-H1*_Hsp_, *ELF3*_Hsp_/*PPD-H1*_Hv_, and *ELF3*_Hsp_/*PPD-H1*_Hsp_, these are also the names with which the lines are referred as (Table 1). However, for the latter combination, no genotype was found without segregation at the *FT1* locus. Despite this limitation, we included the closest available genotype as a representative of the *ELF3*_*Hsp*_*/PPD-H1*_*Hsp*_ combination.

**Table 1.** Genotypic combinations of *ELF3* and *PPD-H1* alleles in the HEB group analysed in this study. The alleles are derived from either the domesticated barley *Hordeum vulgare* (Hv) or the wild progenitor *Hordeum spontaneum* (Hsp) from HEB-25 population.

| Genotype | *ELF3* allele (origin) | *PPD-H1* allele (origin) |
| --- | --- | --- |
| *ELF3*<sub>Hv</sub>/*PPD-H1*<sub>Hv</sub> | Domesticated (*H. vulgare*) | Domesticated (*H. vulgare*) |
| *ELF3*<sub>Hv</sub>/*PPD-H1*<sub>Hsp</sub> | Domesticated (*H. vulgare*) | Wild (*H. spontaneum*) |
| *ELF3*<sub>Hsp</sub>/*PPD-H1*<sub>Hv</sub> | Wild (*H. spontaneum*) | Domesticated (*H. vulgare*) |
| *ELF3*<sub>Hsp</sub>/*PPD-H1*<sub>Hsp</sub> | Wild (*H. spontaneum*) | Wild (*H. spontaneum*) |

The second plant group, names “Bowman group” (Druka et al., 2011) consisted of the wild-type cultivar Bowman and its respective near-isogenic lines (NILs) the allele *elf3* (eam8.w from the line BW290, from Zakhrabekova et al., 2012) and *PhyC-e (*line name BW285, from Pankin et al., 2014). The lines in this group are referred to as BW_WT_, BW_*ELF3*_, and BW_PHYC_, respectively (Table 2). The rationale for having the photoperiod insensitive line BW_*ELF3*_ is to have a line which will express *intrinsic earliness*.

**Table 2.** Genotypic composition of the Bowman introgression group, consisting of the wild-type cultivar Bowman (BW_WT_) and two near-isogenic lines (NILs) differing at the *ELF3* and *PPD-H1* loci: BW_ELF3_, carrying the *elf3* mutant allele (eam8.w), and BW_PHYC_, carrying the *PhyC-e* allele. All lines share the same recessive *ppd-H1* background and differ at either the *ELF3* or *PHYC* locus. Allele origins and mutations are based on Druka et al. (2011), Zakhrabekova et al. (2012), and Pankin et al. (2014).

| Genotype | *ELF3* allele | *PPD-H1* allele | *PHYC* allele |
| --- | --- | --- | --- |
| BW<sub>WT</sub> | *ELF3*<sub>Hsp</sub> | *ppd-H1* | *PhyC-I* |
| BW<sub>ELF3</sub> | *elf3* | *ppd-H1* | *PhyC-I* |
| BW<sub>PHYC</sub> | *ELF3*<sub>Hsp</sub> | *ppd-H1* | *PhyC-e* |

To ensure that the RILs used in the HEB group harboured the different *ELF3* and *PPD-H1* alleles, we sequenced these genomic regions and compared them with Barke and Bowman. Graphical genotyping is provided in Data S1 along with full details of DNA extraction, amplification, and sequencing procedures.

### Experimental design and phenotyping

To ensure precise photoperiod control and minimize light leakage, plants were grown in 80×80×160 cm grow tents (Senua Hydroponics-https://www.senua-hydroponics.com/) under strictly regulated conditions. An illustration of the experimental setup is provided in **Figure S1**. Seeds were sown directly in 0.3 litres pots with Sinclair All Purpose Growing Medium Compost (https://www.sinclairpro.com/). Plants were exposed to 5 photoperiod conditions: 1) 16 h light/8 h darkness, 2) 18 h light/6 h darkness, 3) 20 h light/4 h darkness, 4) 22 h light/2 h darkness and 5) 24 h light/0 h darkness (continuous light). The experiment was repeated independently twice under identical conditions during the spring seasons (March to May) in SRUC’s Peter-Wilson campus (55°55′17.386″ N−3°10′42.175″ E) growth chambers in year 2023 and 2024. Five replicates of each line (7 lines in total) in each condition for each experiment repetition were grown in a completely randomized block design (RBD) within each tent. With 35 pots occupying the 0.64 m^2^ tent space, this resulted in a planting density of 55 plants/m^2^. The light intensity was set at 200 μmol m^−2^ s^−1^ as found to be optimal for promoting robust growth in barley (Yang et al., 2024). Light intensity was measured using a quantum sensor (SKP 200—Skye Instruments www.skyeinstrument.com), while temperature was recorded every 30 minutes using dataloggers (EasyLog USB www.lascarelectronics.com). Temperature was set at 20 °C constant.

Our study concentrated on flowering time response to different photoperiod conditions. The phase duration form emergence to when the awns become visible, defined as heading (BBCH stage 49, Lancashire et al., 1991) was used to determine time to flowering, which was expressed in accumulated thermal time to heading (flowering), assuming a base temperature of 0 °C as described in Parrado et al. (2023).

### Photoperiod Response Modelling

Phenotypic data were manually curated to identify and remove erroneous measurements and biological outliers prior to analysis. Genotype means were visualized across photoperiod conditions, guided by the conceptual framework of Perez-Gianmarco et al. (2019), to select appropriate modelling strategies. This prior knowledge, combined with visual assessment of the data, informed whether: (i) multiple lines could be modelled together while keeping some (or all) photoperiod- response parameters stable, or (ii) the model required inclusion of all parameters (photoperiod sensitivity, threshold photoperiod, and intrinsic earliness) to capture the photoperiod response.

All modelling was performed in RStudio v2.14.4 using the *brms* R-package (Bürkner, 2017).

### Gene expression analysis

To assess whether the observed phenotypic variation could be attributed to the gene of interest rather than background genetic effects in the HEB group, we examined transcript abundance of candidate genes *PPD-H1* and downstream *FT1* in the four lines in the HEB group under contrasting photoperiods: 16h light/8h dark (16h; 20.9°C/16.4°C) and 22h light/2h dark (22h; 19.3°C/15.5°C).

Leaf samples were collected every 6 hours from ZT5 (Zeitgeber Time, indicating the hours after the onset of light) on day 23 post-emergence with two biological replicates and two technical replicates per line, photoperiod condition and time point. Following flash-freezing in liquid N□ and homogenization, total RNA was extracted using Qiagen’s RNeasy Plus Kit with QIAshredder. After DNase treatment (TURBO DNA-free™ Kit) and cDNA synthesis (SuperScript™ III), qPCR was performed in technical duplicate using SYBR® Green chemistry on an AriaMx Real-Time System. *HvTubA* served as the reference gene, showing the most stable expression among tested references (*HvGADPH, HvUbi*).

Expression differences were assessed using two-sided t-tests. Primer sequences and cycling conditions are provided in Table S3.

## Results

To investigate how allelic variation at major flowering time genes affects barley development under different daylengths, we assessed two distinct plant material groups: a set of four recombinant inbred lines of the barley NAM HEB-25 (HEB group) representing combinations of wild and domesticated *ELF3* and *PPD-H1* alleles within a largely isogenic background, and the Bowman near-isogenic line group comprising the wild-type and lines carrying mutant *elf3* or *PhyC-e* alleles. Our primary objective was to characterize the impact of these crucial genes on photoperiod response parameters—including sensitivity, threshold, and intrinsic earliness—across a range of controlled photoperiods. In addition, for the HEB group, we analysed the expression profiles of PPD-H1 and its downstream target FT1 under contrasting photoperiods to examine whether phenotypic differences could be attributed to variation at these flowering time genes. The results below integrate phenotypic modelling and gene expression analysis to reveal how genetic variation at these loci may inform more energy-efficient speed breeding protocols and climate-resilient barley improvement.

### Photoperiod response models on thermal time to heading

To elucidate how genetic variation influences flowering responses under different daylengths, we measured the thermal time to heading across genotypes and modelled photoperiod responses. In the HEB group (Table 1), analysis of mean thermal time to heading for each line (Figure 1a–b) revealed that lines carrying the *PPD-H1*_Hv_ allele exhibited a bi-linear response with a distinct breakpoint at 20 hours. In contrast, lines with the *PPD-H1*_Hsp_ allele showed no response across the photoperiod range studied and consistently flowered earlier under all conditions. Within the *PPD-H1*_Hv_ background, lines with *ELF3*_Hsp_ flowered earlier than those with *ELF3*_Hv_, though both shared similar responsiveness up to the 20-hour breakpoint. In the Bowman group (Table 2), photoperiod sensitivity appeared to be modulated by allelic variation at *PHYC*. Although all genotypes converged to the same flowering time beyond 20 hours, they varied in responsiveness below this threshold.

**Figure 1.**
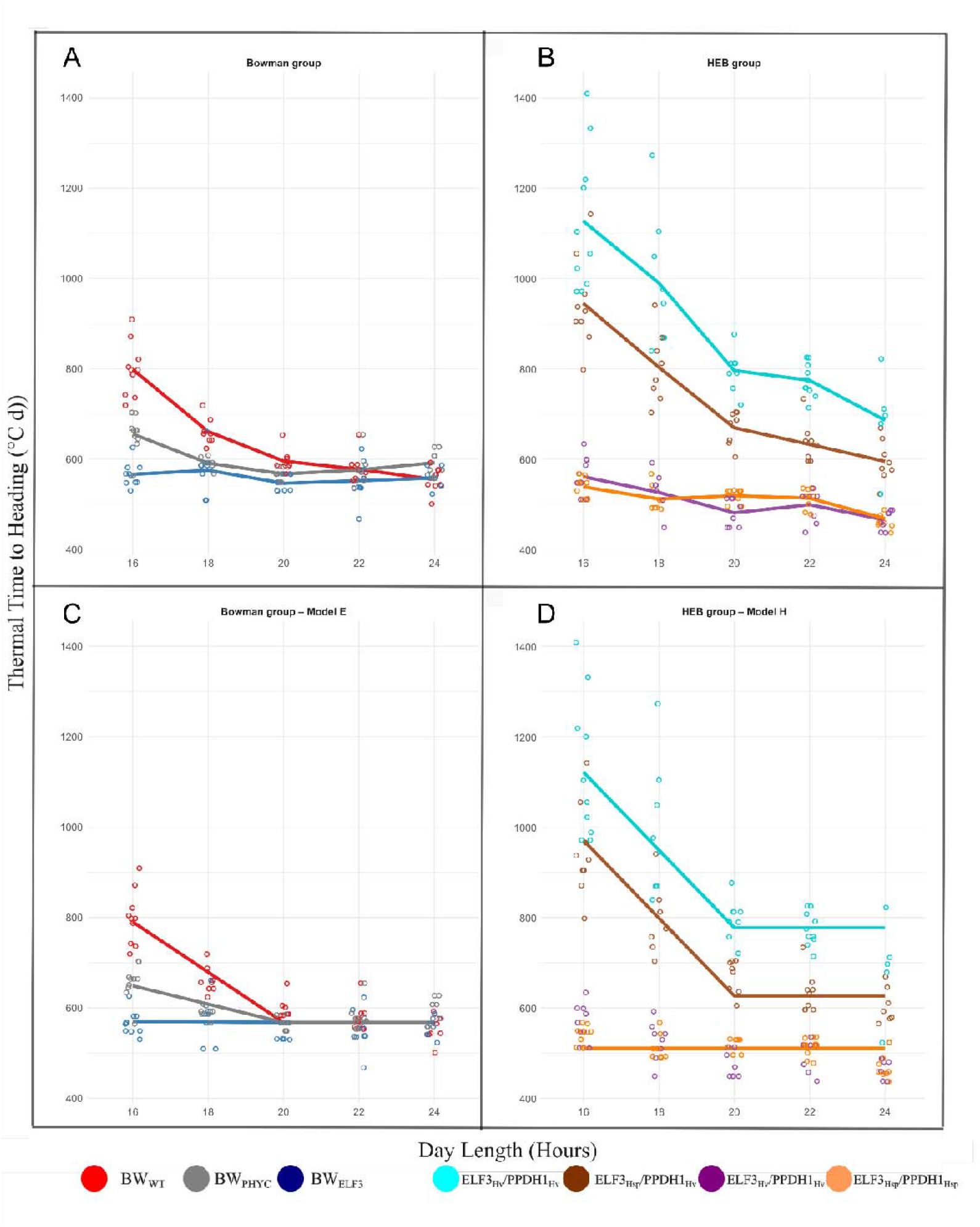
Flowering responses to increasing photoperiods in Bowman and HEB genotypes. (a–b) Observed mean thermal time to heading for each genotype under five photoperiod conditions; (c–d) Fitted Bayesian model based on estimates of photoperiod sensitivity, threshold photoperiod, and intrinsic earliness for each line.

Pairwise Student’s t-tests were conducted to quantify differences in thermal time to heading (GDD) between key photoperiod treatments (16□h vs 20□h and 20□h vs 22□h; Supplementary Table S2a-b). All genotypes except *BW*_*ELF3*_ (as expected) exhibited significant reductions in GDD when the photoperiod was extended from 16□h to 20□h (p□< □0.05), consistent with strong photoperiod sensitivity in this range. However, the magnitude of change was notably smaller in lines carrying the *PPD-H1*_Hsp_ allele compared to lines with photoperiod-responsive alleles (Figure S2). When the photoperiod was further extended from 20□h to 22□h (the photoperiod used in Speed Breeding), did not result in statistically significant changes in GDD in any genotype, indicating convergence of flowering time above the 20□h threshold. These results delineate the photoperiod sensitivity window and highlighted the limited acceleration of *PPD-H1*_*Hsp*_ lines relative to other genotype groups.

These empirical patterns informed our modelling strategy. We first visualized genotype means across photoperiod conditions to determine whether genotypes could be modelled using shared parameters or required separate parameter estimation. We then fitted Bayesian models (Figure 1c–d) to estimate the three key components of photoperiod response: photoperiod sensitivity, threshold photoperiod, and intrinsic earliness. This approach enabled us to identify the specific effects of *PPD-H1, ELF3*, and *PHYC* alleles on the parameters defining our model equations. Detailed model specifications are provided in Supplementary Data S2.

### Gene expression analysis

To further understand the phenotypic differences between HEB lines and photoperiod conditions, we carried out a gene expression analysis via RT-qPCR on *PPD-H1* and *FT1* grown at 16 and 22 h photoperiods. Most of the statistical differences in gene expression between lines were observed at 16h.

*PPD-H1* expression was significantly higher in lines carrying the *PPD-H1*_*Hsp*_ allele compared to those with *PPD-H1*_*Hv*_ at ZT11, ZT17, and ZT23, under both 16 h (Figure 2b) and 22 h (Figure 2d) photoperiods (Student’s t-test, p-values in Supplementary Table S4). Such result is unexpected, since usually *PPD-H1* alleles different at the CCT domain don’t show important differences in expression levels and their effect is usually visible on the expression of downstream genes such as *FT1* (Campoli et al., 2012; Jones et al., 2008; Turner et al., 2005). Notably, at ZT23—the only time point when lights were off in both photoperiod treatments—*PPD-H1*_*Hv*_ expression was undetectable in four out of eight samples in the two lines, correlating with the slower flowering time observed in lines harbouring this allele. At ZT17, *PPD-H1* expression was also significantly higher in *ELF3*_*Hsp*_/*PPD-H1*_*Hv*_ lines compared to *ELF3*_*Hv*_/*PPD-H1*_*Hv*_, correlating to the faster flowering time observed in the first.

**Figure 2.**
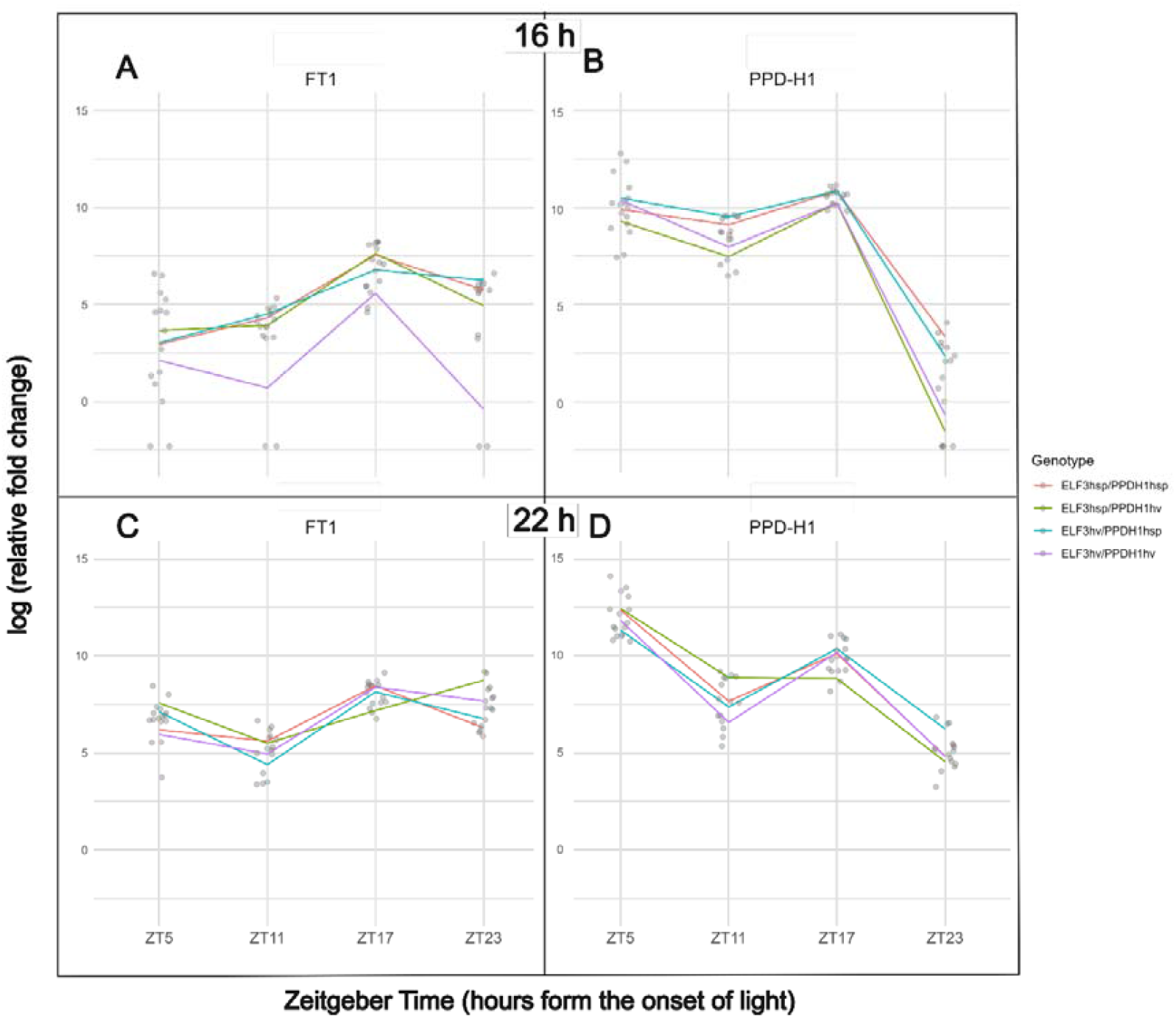
*PPD-H1* and *FT1* expression in four genotypes under two photoperiod conditions (16h light/8h dark and 22h light/2h dark) across four time points (ZT5, ZT11, ZT17, andZT23). Statistical comparisons of mean expression levels (Student’s t-tests) are provided in Tables S4 & S5.

FT1 expression was undetectable in *ELF3*_*Hv*_/*PPD-H1*_*Hv*_ lines at ZT5, ZT11, and ZT23 under the 16 h photoperiod (Figure 2a), with expression absent in two out of four samples at each time point. As expected, statistical differences in *FT1* expression were primarily observed between genotypes differing at the *PPD-H1* locus, with higher expression in lines carrying the *PPD-H1*_*Hsp*_ or *Ppd-H1* allele (Student’s *t*-test, *p*-values in Supplementary Table S5). The only exception was at ZT17 at 16h, where *ELF3*_*Hsp*_/*PPD-H1*_*Hv*_ showed significantly higher expression than *ELF3*_*Hv*_/*PPD-H1*_*Hv*_ despite sharing the same *PPD-H1* allele (Figure 2a). Such result correlates with the faster flowering time observed in *ELF3*_*Hsp*_/*PPD-H1*_*Hv*._

## Discussion

In this study, we build on our previous work (Rossi et al., 2024) by directly testing how different alleles of *PPD-H1, ELF3*, and *PHYC* (*Elf3* or *ELF3*_*Hv*_, *elf3, ELF3*_*Hsp*_, *ppd-H1* or *PPD-H1*_*Hv*_, *Ppd-H1* or *PPD-H1*_*Hsp*_, *PhyC-I*, and *PhyC-e*) affect key aspects of photoperiod response in barley under long days (16-18h) and very long days (above 18h) in controlled environments. To achieve this, we used both near isogenic lines in the Bowman background recombinant inbred lines from the HEB-25 NAM population, selected to minimize genetic variation outside the target loci. This approach enables us to directly quantify the effects of specific alleles on key aspect of the photoperiod response.

Our current study is the first to assess how allelic variation at three major flowering time genes collectively influence threshold photoperiod, photoperiod sensitivity, and intrinsic earliness (Major et al., 1980, Perez-Gianmarco et al., 2019) within a single, unified experiment. By integrating flowering time data across a range of day lengths with gene expression analysis, we directly compared the effects of distinct allelic combinations at these loci on photoperiod response parameters. This comprehensive approach yields new insights into the genetic control of flowering in barley—specifically, by identifying the photoperiod threshold necessary to optimize energy-efficient speed breeding protocols (Watson et al., 2018; Hickey et al., 2019) and by elucidating patterns of photoperiod sensitivity and intrinsic earliness that will guide allele selection for adaptation to a warming climate (Slafer & Rawson, 1995; Craufurd et al., 2009; Reynolds et al., 2009).

Although our results revealed some statistically significant differences in thermal time to heading between long photoperiod treatments (such as 16□h compared to 20□h in lines harbouring *Ppd-H1*), the magnitude of these differences was limited. Accordingly, we have focused our interpretation on the photoperiod response model, since it better captures the underlying biological mechanisms and supports robust comparisons across different genotypes and environments. Our findings show that, lines carrying the *ppd-*H1 allele, including *ELF3*_*Hv*_*/PPD-H1*_*Hv*_, *ELF3*_*Hsp*_*/PPD-H1*_*Hv*_, *BW*_*ELF3*_ and *BW*_*PHYC*_—regardless of *ELF3* background—exhibited a clear bi-linear response to photoperiod, with flowering time accelerating below a threshold of 20 hours and then plateauing. In contrast, lines with the *Ppd-H1* allele, including *ELF3*_*Hv*_*/PPD-H1*_*Hsp*_ and *ELF3*_*Hsp*_*/PPD-H1*_*Hsp*_, showed no substantial response to increasing photoperiod and flowered early under all conditions, indicating that lines harbouring this allele had already reached the threshold photoperiod at 16h. Consistent with the early-flowering phenotype, *PPD-H1* expression was significantly higher in lines carrying the *PPD-H1*_*Hsp*_ allele compared to those with *PPD-H1*_*Hv*_ under 16h. This result is unexpected, as previous studies have reported that *PPD-H1* alleles differing at the CCT domain typically do not show substantial differences in expression levels (Turner et al., 2005; Jones et al., 2008; Campoli et al., 2012). Such differences in expression correlate with levels of *FT1* expression with significant higher expression on lines carrying the *Ppd- H1*, In contrast to Parrado et al., (2023) different *PPD-H1* alleles yielded a different intrinsic earliness and this response correlated with a significant higher level of expression in *Ppd-H1* lines at *PPD-H1* and *FT1* in 22 h (after the threshold photoperiod). In addition, *ELF3*_*Hsp*_ lines consistently headed earlier than *ELF3*_*Hv*_ in a *ppd-H1* background, indicating that functional *ELF3* alleles primarily shift intrinsic earliness rather than threshold or sensitivity, this correlated with a significant higher expression of *FT1* in *ELF3*_*Hsp*_/*PPD-H1*_*Hv*_ than *ELF3*_*Hv*_/*PPD-H1*_*Hv*_ at ZT17 in 16h. Whereas the *Phyc-e* allele, harboured within the Bowman NIL *BW*_*PHYC*_, affected photoperiod sensitivity but not intrinsic earliness.

As climate warming alters the growing season in parts of Europe, the adaptive advantage conferred by *ppd-H1* alleles in barley cultivars is likely to diminish, highlighting the need to address this issue in future breeding strategies (Herzig et al., 2018). Our analysis presented in this paper helps define how specific allelic combinations influence photoperiod sensitivity and intrinsic earliness and indicates that combining *Phyc-e* or *ELF3*_*Hsp*_ alleles in a *ppd- H1* background offers a valuable alternative to using *Ppd-H1* alleles. Our results demonstrate that specific allele combinations enable intermediate flowering times and retain photoperiod sensitivity, providing greater flexibility for breeding climate-resilient varieties. By facilitating crops to escape heat and drought stress without excessively shortening the growing season, this strategy could offer a more balanced adaptation to warming climates than relying exclusively on *Ppd-H1*. In addition, the *ppd-H1* allele has been shown to enhance spikelet survival by reducing tip degeneration (Huang & Schnurbusch, 2024) and to promote greater floral primordia survival, which translates into improved spike fertility and potentially higher yield in both *Phyc-e* and *Phyc-i* background (Parrado et al., 2025). Furthermore, *ELF3*_*Hsp*_ has recently been discovered to contribute to phenotypic and developmental acclimation to elevated temperatures (Zhu et al., 2023). Taken together, these findings support the continued exploration and strategic deployment of of *ppd-H1* alleles, especially in combination with alleles such as *Phyc-e* or *ELF3*_*Hsp*_.

The data and observations obtained from this study provide valuable practical insights: the threshold photoperiod help us understand how to optimize genotype-tailored speed breeding protocols. Current speed breeding approaches for long-day crops often maintain an extended 22-hour photoperiod with a two-hour dark phase and cooler nights to accelerate generation turnover and mitigate stress (Watson, 2019). Many major crops, including wheat, barley, canola, chickpea, pea, durum wheat, and oat, are routinely grown under a 22-hour photoperiod as the accepted speed breeding protocol (Watson et al., 2018; González-Barrios et al., 2021). Our results indicate, however, that photoperiod requirements for optimal flowering vary considerably depending on the alleles present at the *PPD-H1* locus. Lines with the *ppd-H1* allele reach a threshold photoperiod at 20 hours, while those with the *Ppd- H1* allele are unresponsive to photoperiods longer than 16 hours—a pattern also observed by Parrado et al. (2023). This suggests that photoperiod length can be fine-tuned to the genetic background of breeding materials: for all genotypes, the widely used 22-hour light regime exceeds what is necessary for accelerated flowering. Based on calculations from Zhang et al. (2017) and Teo et al. (2024), as described in the introduction, adjusting photoperiods accordingly could reduce lighting demands I the range of 9%–27%—or 4.5–13.5 kWh per 8 sq. ft. annually—translating to both energy and cost savings across facilities. At current UK commercial electricity rates (approximately £0.22 per kWh; UK Department for Energy Security and Net Zero, 2024), these savings would equate to £1.240 to £3.715 for a 10.000 sq. ft. commercial greenhouse for a 16h and a 20h photoperiod, respectively, compared to a 22h photoperiod. Moreover, reducing lighting hours not only lowers direct electricity costs, but also decreases the demand for cooling and ventilation, further enhancing the sustainability and cost-efficiency of breeding facilities.

NILs are invaluable tools for dissecting the effects of individual loci on complex traits such as flowering time and photoperiod sensitivity, enabling precise attribution of phenotypic variation to specific genetic changes. Developing NILs that carry distinct *ELF3* and *PPD-H1* wild and domesticated alleles would be an important resource for future studies aiming to optimize barley ideotypes for climate adaptation. Furthermore, evaluating these key allelic combinations—including variants at *PHYC*—in field experiments across different sowing dates will be essential for validating their performance and stability under variable seasonal and environmental conditions.

## Supplementary data

**Tables S1–S5:** Supplementary tables containing additional data referenced in the main text.

**Table S1**. Genotype data from the Infinium iSelect 50k SNP chip of HEB group lines for the seven major flowering time loci. The markers utilized for flowering time loci were sourced from the panel of pre-selected (SNPs) described by Zahn et al. (2023)

**Table S2a–b**. The significance of differences in GDD to heading between photoperiod treatments (16 h vs. 20 h and 20 h vs. 22 h) was determined for each genotype by Student’s t-test, with mean ± SD reported and a significance level of p < 0.05.

**Table S3**.

(a) List of primer pairs used in the resequencing of the *PPDH1* and *ELF3* genomic regions.

(b) List of primers employed in Sanger sequencing; primers in bold provided acceptable sequences for the haplotype analysis.

(c) List of primer pairs for gene expression analysis.

**Table S4**. Significant differences in PPD-H1 expression fold-change rates between genotypes of HEB lines group within the same photoperiod condition, quantified via RT-qPCR and assesed using Student’s t-test at different time points

**Table S5**. Significant differences in FT1 expression fold-change rates between genotypes of HEB lines group within the same photoperiod condition, quantified via RT-qPCR and assesed using Student’s t-test at different time points

**Data S1:** Molecular characterization of *ELF3* and *PPD-H1* allelic variation in HEB lines.

**Data S2:** Overview of the photoperiod response modeling strategy.

**Figure S1.**
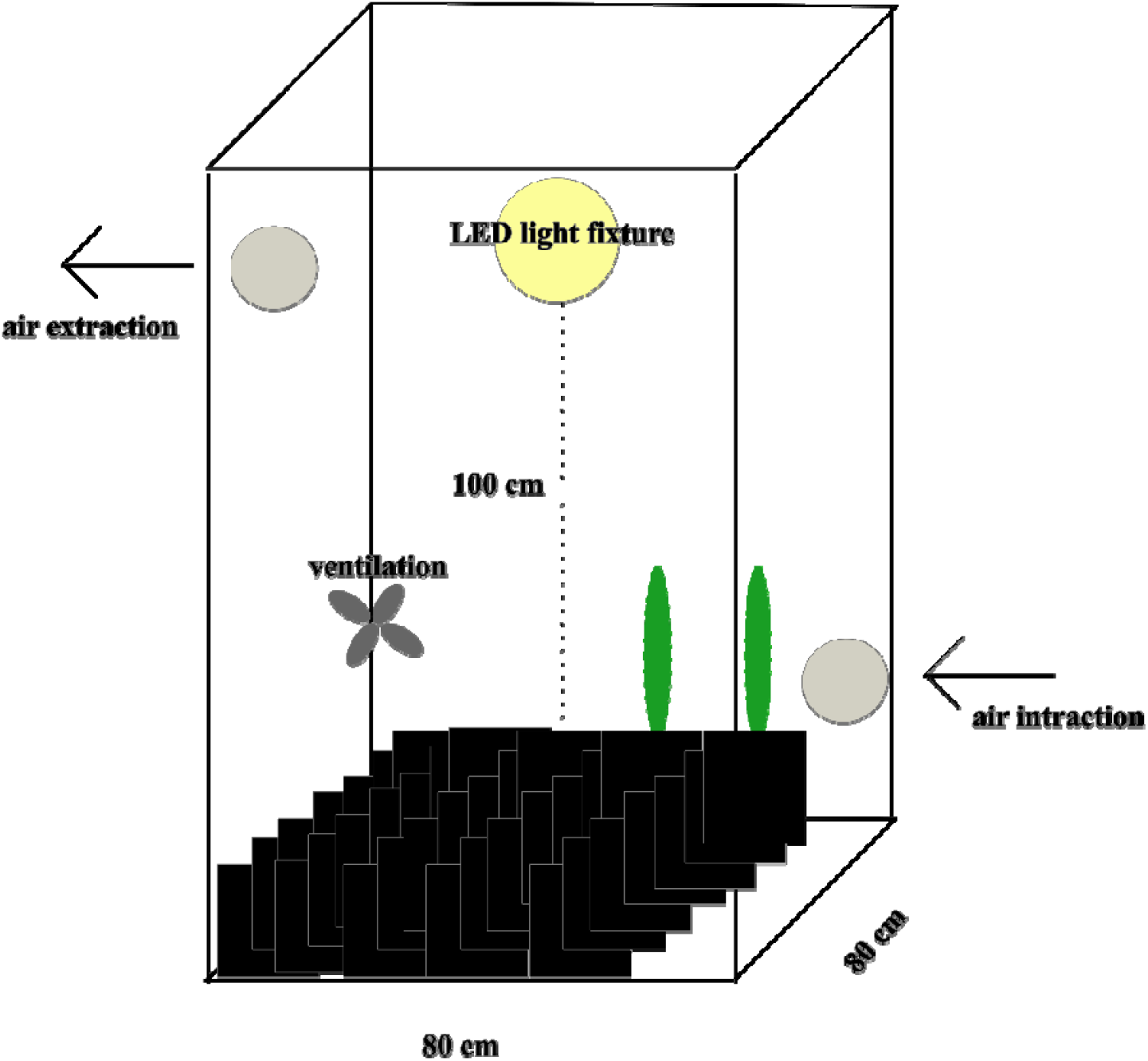
Illustration of the grow tent setup showing how different light regimes were applied across experimental conditions in the photoperiod study.

**Figure S2.**
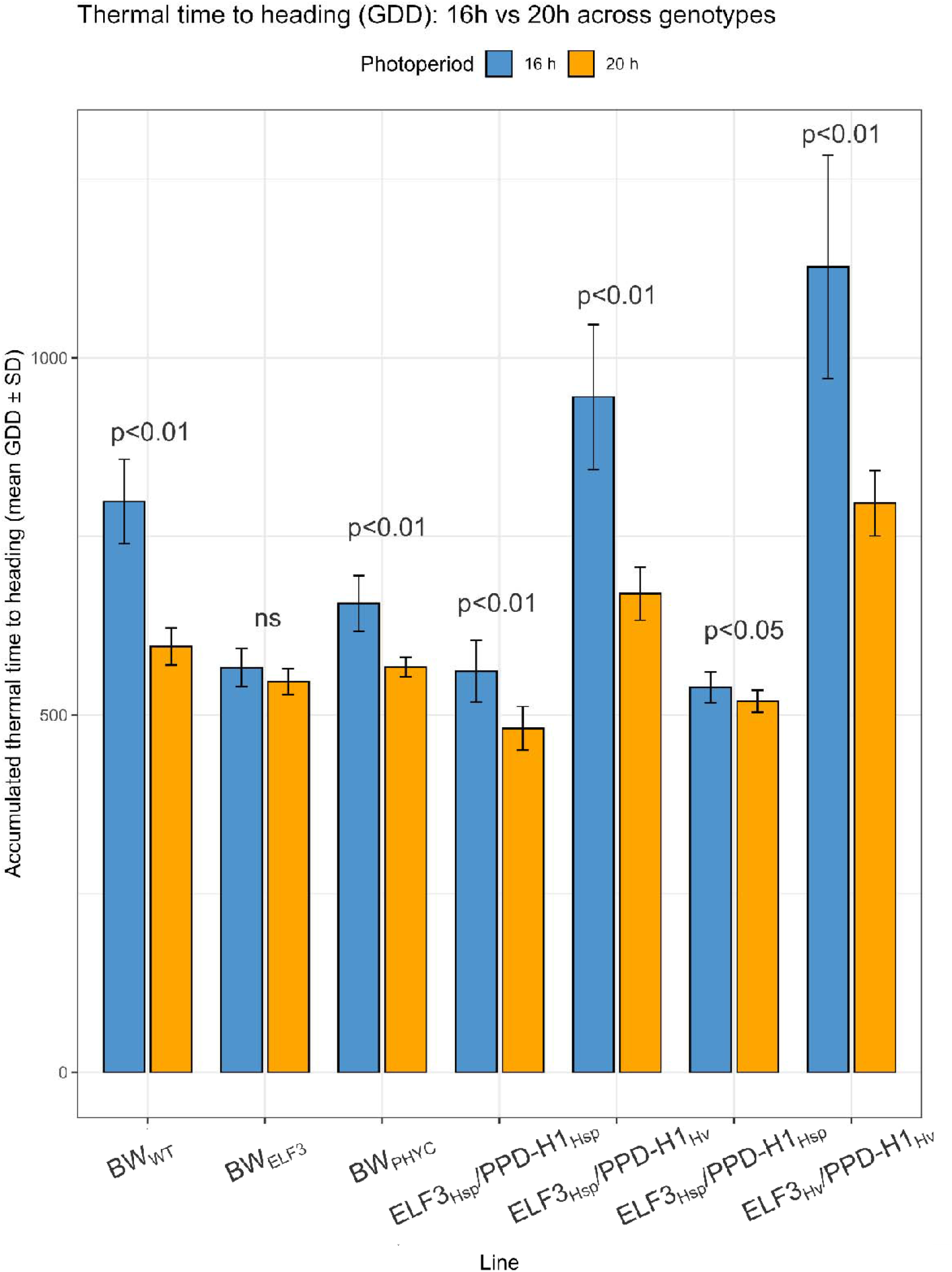
Barplots of thermal time to heading under 16□h and 20□h photoperiods across lines.

## Acknowledgements

This project was funded by EASTBIO DTP and BBSRC to Nicola Rossi and by direct funding to Rajiv Sharma from SRUC. We thank Gustavo Slafer for helpful discussion throughout the work. We would also like to acknowledge Neil Havis, Kalina Gorniak, Julie Fortune, Grace Cuthill, and Lachlan Jones from the crop and soils department-SRUC for their technical help in conducting the controlled environments experiments.

The authors have no relevant financial or non-financial interests to disclose.

## Author contributions

Nicola Rossi performed the investigation, formal analysis, and preparation of the original draft. Conceptualization, methodology, and funding acquisition were carried out jointly by Nicola Rossi, Rajiv Sharma, Wayne Powell, and Karen Halliday. All authors contributed to the review and editing of the manuscript.

## Data availability

Data will be made available on request.

